# A Targeted Vaccine against COVID-19: S1-Fc Vaccine Targeting the Antigen-Presenting Cell Compartment Elicits Protection against SARS-CoV-2 Infection

**DOI:** 10.1101/2020.06.29.178616

**Authors:** Andreas Herrmann, Junki Maruyama, Chanyu Yue, Christoph Lahtz, Heyue Zhou, Lisa Kerwin, Whenzong Guo, Yanliang Zhang, William Soo Hoo, Soonpin Yei, Sunkuk Kwon, Yanwen Fu, Sachi Johnson, Arthur Ledesma, Yiran Zhou, Yingcong Zhuang, Elena Yei, Tomasz Adamus, Slobodan Praessler, Henry Ji

## Abstract

Vaccination efficacy is enhanced by targeting the antigen-presenting cell compartment. Here, we show that S1-Fc antigen delivery targeting the FcγR^+^ antigen-presenting cell compartment elicits anti-SARS-CoV-2 S1-antigen specific IgG production *in vivo* exerting biologically functional and protective activity against live virus infection, assessed in a stringent experimental virus challenge assay *in vitro*. The S1-domain of the SARS-CoV-2 spike protein was genetically fused to a human immunoglobulin Fc moiety, which contributes to mediate S1-Fc cellular internalization by FcγR^+^ antigen-presenting cells. Immediately upon administration intramuscularly, our novel vaccine candidate recombinant rS1-Fc homes to lymph nodes *in vivo* where FcγR^+^ antigen-presenting cells reside. Seroconversion is achieved as early as day 7, mounting considerably increased levels of anti-S1 IgGs *in vivo*. Interestingly, immunization at elevated doses with non-expiring S1-Fc encoding dsDNA favors the education of a desired antigen-specific adaptive T cell response. However, low-dose immunization, safeguarding patient safety, using recombinant rS1-Fc, elicits a considerably elevated protection amplitude against live SARS-CoV-2 infection. Our promising findings on rS1-Fc protein immunization prompted us to further develop an affordable and safe product for delivery to our communities in need for COVID-19 vaccinations.

## Introduction

The pandemic Coronavirus Disease 2019 (COVID-19), induced by Severe-Acute-Respiratory-Syndrome Coronavirus-2 (SARS-CoV-2) infection, was declared a Public Health Emergency of International Concern in January 2020^1^. In March 2020, more than 80,000 infections and 3,000 deaths were reported in China. COVID-19 became pandemic by spreading to numerous countries including the United States, Germany, Italy, Spain, Iran, Japan, Korea and more. By April 2020, the global infection rate was reported at 870,000 with 43,000 deaths^2^. In June 2020, approximately 10 million infection were estimated with 500,000 deaths caused by COVID-19^3^.

SARS-CoV-2 belongs to the virus family *Coronaviridae* (betacoronavirus taxon, or type II coronaviruses)^4^ and is commonly known to be pathogenic to humans upon infection^5^. Alphacoronavirus and most betacoronavirus infections manifest with symptoms similar to the common cold^6^. However, SARS-CoV-2 infection exerts an elevated pathology, manifesting as a severe atypical pneumonia, fever, acute respiratory distress syndrome (ARDS), sepsis and potentially patient death.

SARS-CoV-1 was initially identified and characterized as a novel coronavirus causing ARDS in 2003^7-9^. Efforts have been made to develop therapeutic and prophylactic interventions. However, all efforts were retired in 2007 when the virus was eradicated from the human population by non-pharmaceutical interventions^2^. None of the vaccines had gone past clinical testing phase I^2^. Currently, therapeutics and vaccines against SARS-CoV-2 are not available but are critically needed.

Since the initial discovery of SARS-CoV-1 in 2003, a repertoire of vaccination approaches underwent preclinical and clinical testing, ranging from attenuated virus to antigen-encoding nucleic acid sequences, differing in immunogen composition, administration format and route^2, 10^. Pioneering antigen-specific vaccination studies identified and employed the spike protein, which was administered encoded by nucleic acid sequence, to serve as a promising immunizing antigen^11^. The spike transmembrane glycoprotein, an immunologically unique integral SARS-CoV-1 and SARS-CoV-2 protein, consists of an extracellularly exposed N-terminal S1 domain (including a receptor binding domain, RBD) engaging with host receptor ACE2 in humans^12^, followed by a S2 domain mediating virion: host-cell membrane fusion resulting in syncytia formation^12^, which is associated with aggressive disease progression^5, 9^. A transmembrane region and an intracellular carboxyterminal cytoplasmic domain complete the spike protein^12^. In 2004, Nabel and colleagues reported a promising plasmid DNA vaccine candidate against SARS-CoV-1 tested in mice, introducing the extracellular domains S1 and S2 as well as the transmembrane domain as immunizing antigen^11^. In 2020, an advanced vaccine candidate employed inactivated SARS-CoV-2 virus acquiring immunization efficacy^13^. Both studies assess promising humoral as well as adaptive immune responses, which are considered to contribute to successful vaccination and long-term protection. While the primary goal of immunization is the production of antigen-specific antibodies capable of virus neutralization and eliciting protective activity^14^, the antigen-specific education of the T cell compartment is thought to mount an antiviral response by interferon (IFN) production^15^. Furthermore, CD4^+^IFNγ^+^ Th1 polarization and maturation of virus-antigen specific CD8^+^ T cells contribute to enhanced anti-viral adaptive immune responses. Interestingly, interleukin-6 (IL-6) secretion has been found to be associated with advanced pulmonary pathology in COVID-19 patients^16^, which draws particular attention to IL-6 circuitry-counteracting IFN responses^17^ and the importance of an operative adaptive immune response. However, directing an immunization approach against COVID-19 to directly target the antigen-presenting cell (APC) compartment, which has been emphasized by Steinman and colleagues to dramatically enhance both, the humoral and adaptive immune response^18^, remains elusive.

Here, we report a unique targeted immunization approach against COVID-19, directly delivering the immunizing S1-Fc antigen to FcγR^+^ APCs. Our immunization approach focusses on the SARS-CoV-2 spike protein S1 domain serving as the viral antigen. We genetically fused the S1 domain to the Fc moiety of human immunoglobulin IgG1, which facilitates cellular internalization of S1-Fc by FcγR^+^ APCs, such as dendritic cells and macrophages as well as B cells, where IgG1-Fc is known to crossreact with murine FcγR^19^. Targeted immunization with rS1-Fc protein potentially affords low dose and therefore safe immunization, associated with enhanced desired immune responses^18^. Immediately after rS1-Fc administration, rS1-Fc homes to the inguinal lymph node where FcγR^+^ APCs reside. Furthermore, rS1-Fc immunization induces rapid seroconversion and mounts a considerably elevated production of S1-specific antibodies *in vivo*, indicating a robust S1-Fc-induced humoral immune response. Interestingly, persistent delivery of S1-Fc by immunization with non-expiring S1-Fc dsDNA induces a potent adaptive immune response, including immunization-dose-dependent Th1 polarization and CD8^+^IFNγ^+^ effector T cell maturation *in vivo*. Moreover, high-dose administration of non-expiring S1-Fc dsDNA induces the education of anti-S1 antigen-specific CD8^+^ T cells. However, in direct comparison, immunization with low-dose recombinant rS1-Fc protein induces the production of S1-specific antibodies, exerting considerably elevated neutralization activity against SARS-CoV-2 virus *in vitro*. Hence, our promising findings prompted us to initiate further proof-of-concept/efficacy studies as well as toxicologic studies, employing rS1-Fc immunization followed by live SARS-Cov-2 virus challenge, to further develop an affordable, readily scalable manufacturing and safe product for delivery to our communities in need for COVID-19 vaccinations.

## Results

The S1-Fc fusion protein serves as the antigen to achieve immunization. The human immunoglobulin Fc moiety is thought to engage with FcγR^+^ expressing cells such as dendritic cells, macrophages and B cells, thus, targeting the antigen-presenting cell compartment^20^. The S1 domain of the SARS-CoV-2 spike protein serves as the immunogenic antigen to mount anti-viral humoral and adaptive immune responses. Upon production of a linear S1-Fc encoding dsDNA, harboring a constitutively active promoter and a cell secretion sequence (Fig. 1A), we assessed S1-Fc protein secretion by murine muscle cells upon delivery. Linear S1-Fc encoding dsDNA was internalized immediately upon electroporation by murine fibroblasts and muscle cells with considerably high efficacy as shown by flow cytometry (Fig. 1B), which could be validated by confocal microscopy (Fig. 1C). Subsequently, critical secretion of S1-Fc protein, which is thought to serve as the immunizing antigen, was detected in muscle cell culture supernatant (Fig. 1D). Notably, S1-Fc protein was detected in the *biceps femoris* muscle 44 days after administration, which is indicative of continuous non-exhausting S1-Fc protein *de novo* synthesis and S1-Fc-protein release (Supplemental Fig. 1A). Moreover, S1-Fc protein was found internalized by dendritic cell species in *biceps femoris*, suggesting operative targeting of the FcγR^+^ APC compartment (Supplemental Fig. 1B).

**Figure 1:**
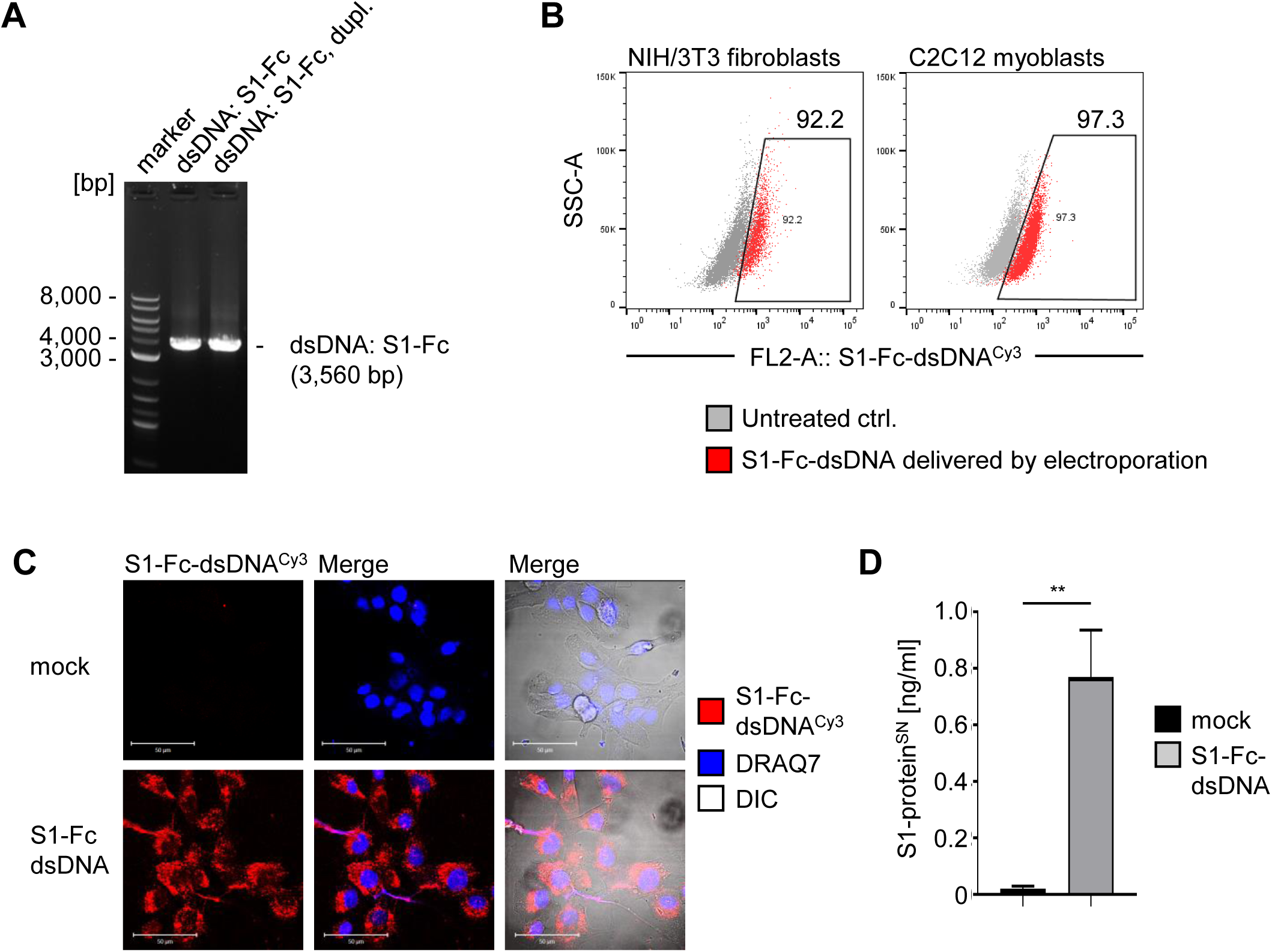
Production and secretion of S1-Fc-protein antigen by muscle cells. (**A**) Linear dsDNA encoding S1-Fc was generated by PCR-based amplification and subjected to analytical DNA gel electrophoresis, assessing accurate size and excluding undesired intermediate product. (**B**) Cellular internalization of fluorescently labeled dsDNA encoding S1-Fc was assessed by flow cytometry immediately after electroporation procedure, gating on dsDNA encoding S1-Fc^+^ cells (red). (**C**) Cellular internalization of S1-Fc-dsDNA was validated by confocal microscopy. Scale, 50 μm. (**D**) Produced S1-Fc protein secreted by C2C12 muscle cells within 96 h after electroporation was assessed from collected cell supernatant and analyzed by MSD assay. SD shown. T-test: **) *P* < 0.01.

Recombinant rS1-Fc protein chimera was produced in CHO cells and subjected to analytical size-exclusion chromatography showing a distinct elution of rS1-Fc monomer (84.5%). However, a minor molecular population (13.2%) eluted as self-assembled (rS1-Fc)_n_ aggregate (Supplemental Fig. 2A) which might by owed to intrinsic oligomerization activity of the SARS-CoV-2 spike protein^21^. Notably, gel electrophoretic analysis showed reduced rS1-Fc aggregation activity in repeated protein production (Supplemental Fig. 2B). rS1-Fc immunologic identity was validated by immunodetection of S1 and Fc upon Western blotting procedure (Fig. 2A). Chimeric rS1-Fc is readily internalized by murine RAW264.7 macrophages *in vitro* (Fig. 2B) and can be found throughout the cell cytoplasm as well as the cell nucleus as demonstrated by confocal laser-scanning microscopy (Fig. 2C). Importantly, rS1-Fc engages with ACE2 as assessed by ELISA (Fig. 2D). The S1 domain of the SARS-CoV-2 spike protein binding to host receptor ACE2 was emphasized to contribute critically to the initiation of host cell infection^12^. However, rS1-Fc binding to ACE2 indicates operative functionality, and furthermore, binding represents a decoy activity by latent competition with viral spike protein for ACE2 binding, potentially reducing host cell virus susceptibility. Moreover, targeted delivery of rS1-Fc to the APC compartment is thought to be mediated by Fc:FcγR interaction, which requires rS1-Fc homing to secondary lymphoid organs once administered. Intramuscular administration into the *biceps femoris* facilitates rS1-Fc migration to the inguinal lymph node within 1 hour as demonstrated by longitudinal nIR fluorescence imaging (Fig. 2E, F). Homing to the lymph node critically enhances the exposure of rS1-Fc to FcγR expressing APCs and favors its cellular internalization and intended processing. Interestingly, S1-protein^+^ splenic APCs were observed 44 days after initial intramuscular administration of rS1-Fc (Fig. 2G).

**Figure 2:**
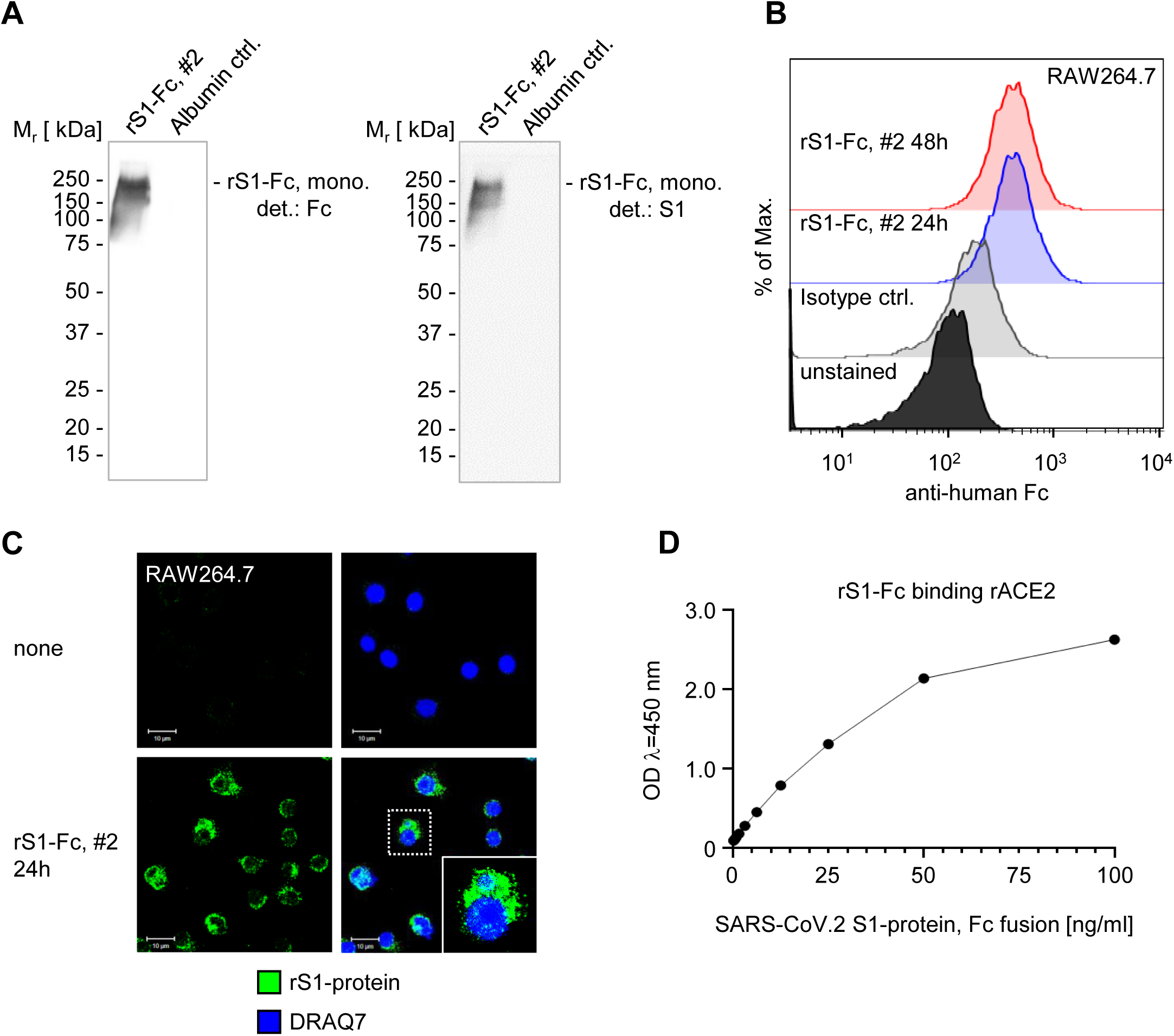
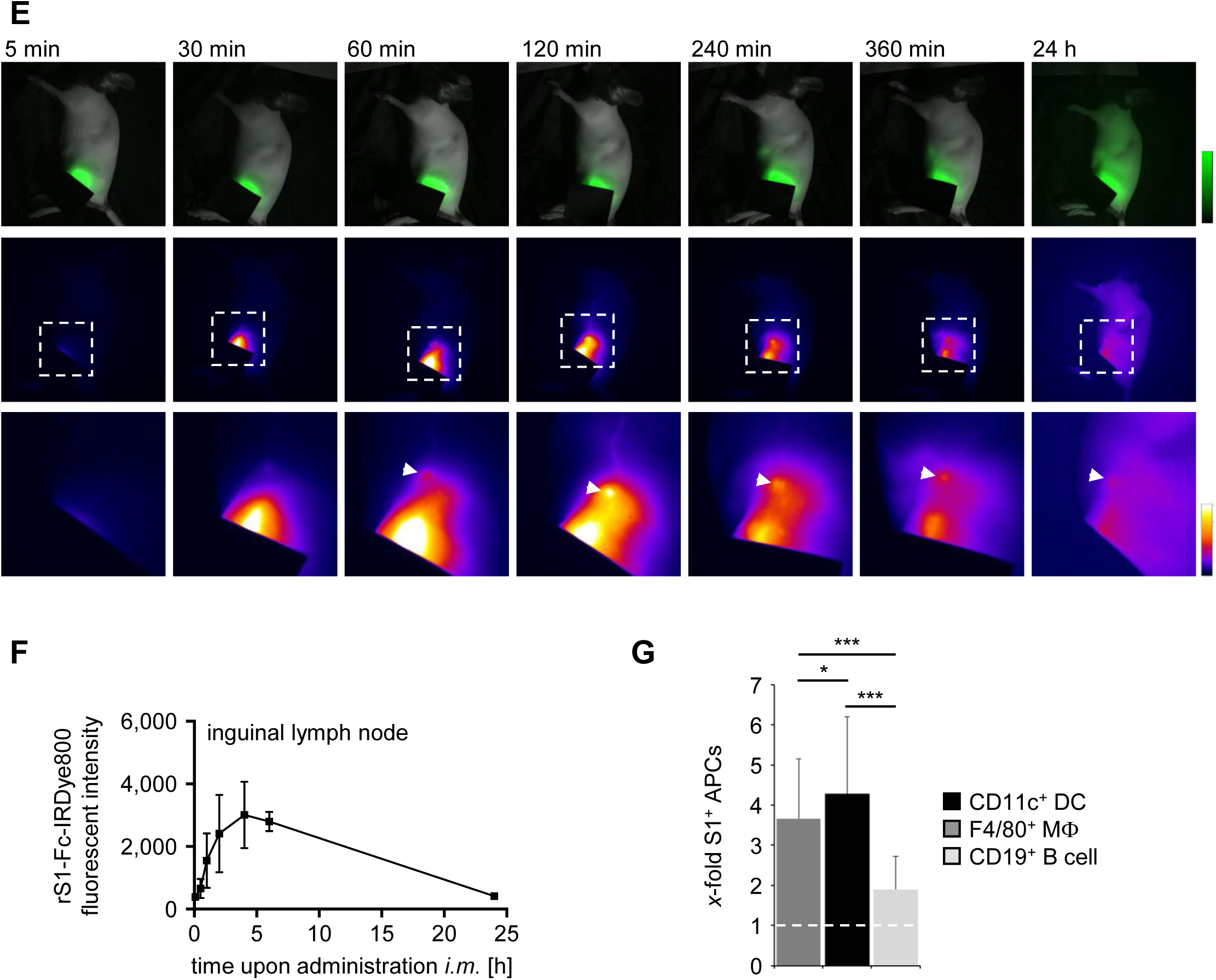
Generation of functional rS1-Fc-prtoein antigen. Recombinant rS1-Fc protein was characterized by SEC HPLC and subsequently (**A**) subjected to Western blot analysis immunologically detecting and confirming human Fc (left) and SARS-CoV-2-S1-domain (right) migrating identically. (**B**) Murine macrophages RAW264.7 internalize rS1-Fc as assessed by flowcytometric analysis and (**C**) confirmed by confocal microscopy. Scale, 10 μm. (**D**) Generated rS1-Fc is functionally binding to human ACE2 as shown by ELISA. (**E, F**) rS1-Fc homes to the inguinal lymph node within an hour upon administration into the *biceps femoris* as determined by longitudinal nIR imaging of the lymphatics. SD shown. (**G**) rS1-Fc and/or S1 and/or processed S1 peptide was found 44 d upon initial administration in splenic APCs as assessed by flow cytometry.

Freshly isolated splenic APCs, such as CD11c^+^ dendritic cells, F4/80^+^ macrophages and CD19^+^ B cells, readily internalize rS1-Fc within 30 min as assessed by flow cytometric analysis (Fig. 3A). Notably, prolonged exposure of splenic APCs to rS1-Fc did not result in increased cellular load, which is suggesting a cytologic limitation in antigen uptake. However, blocking CD16/32^+^ FcγR resulted in considerably reduced rS1-Fc cellular internalization, indicating functional targeting of rS1-Fc into the FcγR^+^ APC compartment (Fig. 3B).

**Figure 3:**
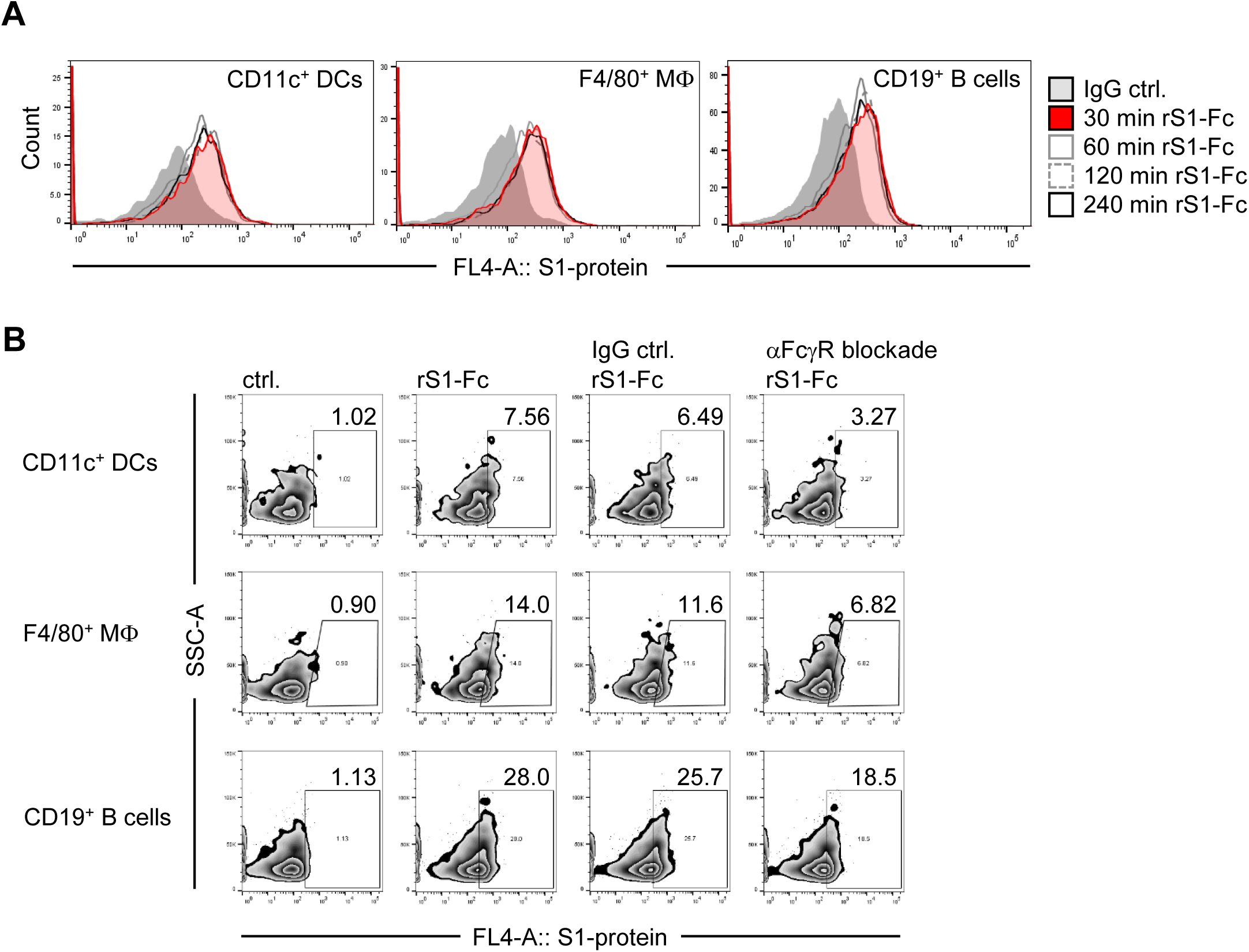
Fc moiety of S1-Fc facilitates functional targeting into the FcγR^+^ APC compartment. (**A**) Splenic cell populations were exposed to rS1-Fc for time points as indicated and rS1-Fc cellular internalization by CD11c^+^ dendritic cells, F4/80^+^ macrophages and CD19^+^ B cells was assessed by flow cytometry. (**B**) Reduced rS1-Fc cellular internalization was determined by exposing splenic cell populations to rS1-Fc with or without CD16/CD32 FcγR^+^ blocking antibody as indicated and S1-protein^+^ cell populations were gated on.

Immunization of mice with either linearized dsDNA encoding S1-Fc or recombinant rS1-Fc protein was administered into the *biceps femoris* intramuscularly. Repeated “boosting” immunization was scheduled at day 21 after initial immunization administration. Blood serum of immunized mice was collected weekly to facilitate monitoring of anticipated immune responses (Fig. 4A). Immunization of mice with linearized dsDNA encoding S1-Fc mounted a significant and robust CD4^+^IFNγ^+^ Th1 polarization *in vivo* in a dose-dependent manner (Fig. 4B). Moreover, S1-antigen specific CD8^+^ T cells isolated from spleen accumulated upon immunization at increased dose (Fig. 4C). Furthermore, high dose immunization favored CD8^+^IFNγ^+^ effector T cell *in vivo* education in a dose-dependent manner (Fig. 4D). Thus, dose-dependent adaptive immune responses upon administration of non-expiring S1-Fc dsDNA indicate, that considerably elevated dosing with non-expiring S1-Fc dsDNA is required to elicit a desired adaptive T cell immune response, continuously feeding S1-antigen systemically. In turn, the humoral B cell immune response is sensitive to reduced immunization doses. Complete seroconversion is detectable at day 10 upon initial immunization with both, 50 μg and 20 μg of administered non-expiring S1-Fc dsDNA, mounting similar levels of S1-specific serum IgG antibodies. However, administration of 2 μg non-expiring S1-Fc dsDNA did not suffice to induce a robust production of S1-specific serum IgG antibodies (Fig. 4E). Remarkably, immunization with recombinant rS1-Fc facilitated accelerated seroconversion detectable at day 7, mounting considerably increased levels of S1-specific serum IgG antibodies. Production of S1-specific serum IgG antibodies *in vivo* experiences a decrease at day 24, which might be owed to the anticipated feedback inhibition loop mediated by FCγR-B activity, facilitating self-regulatory termination of antibody production^22-25^. However, upon administration of a repeated “boosting” immunization at day 21, S1-specific serum IgG antibody production recovered by day 28 resulting in considerably elevated and higher anti-S1 IgG serum levels (Fig. 4F). Moreover, murine blood serum seropositive for anti-S1 IgG significantly reduced the interaction of the viral S1-domain and host receptor ACE2 at high dilution, indicating production of a highly potent anti-S1 IgG serum antibody population induced by rS1-Fc protein immunization (Fig. 4G). Importantly, collected blood serum seropositive for anti-S1 IgG elicits protection against live SARS-CoV-2 infection in a stringent experimental virus challenge assay. However, although continuous S1-Fc expression and antigen production is facilitated by immunization with non-expiring S1-Fc-encoding dsDNA, we observed that high dose immunization with non-expiring S1-Fc-encoding dsDNA elicits protection up to 50% only. In contrast, low-dose immunization with recombinant rS1-Fc administered intramuscularly mounted considerable increased protection activity (80%) (Fig. 4H). Interestingly, routing rS1-Fc administration via intravenous injection resulted in a similar protection efficacy (77.7%), indicating that rS1-Fc immunization propagates a robust humoral immune response. Hence, immunization against SARS-CoV-2 employing rS1-Fc protein unfolds rapid seroconversion and production of anti-S1-specific IgG eliciting protection capacity.

**Figure 4:**
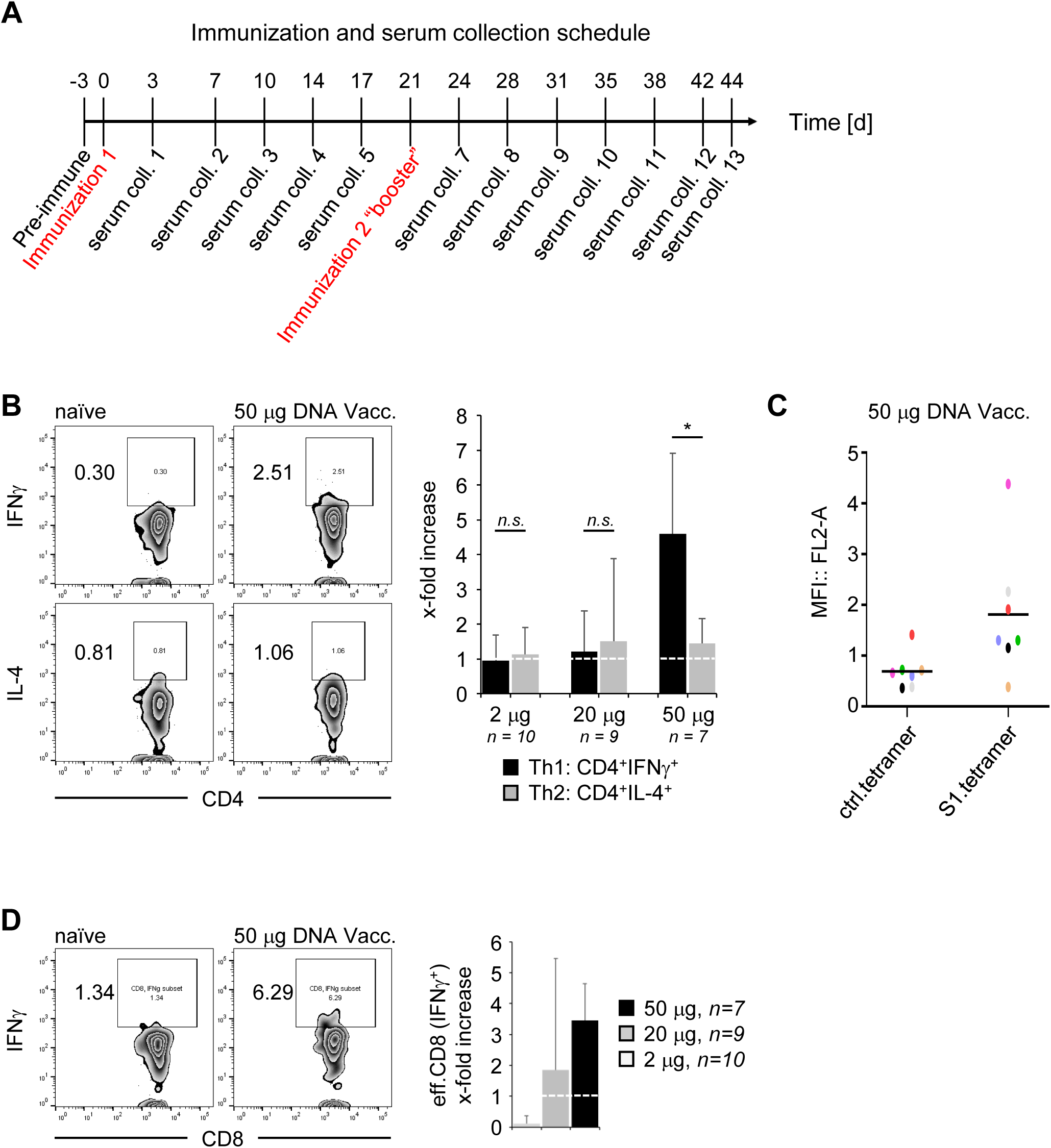
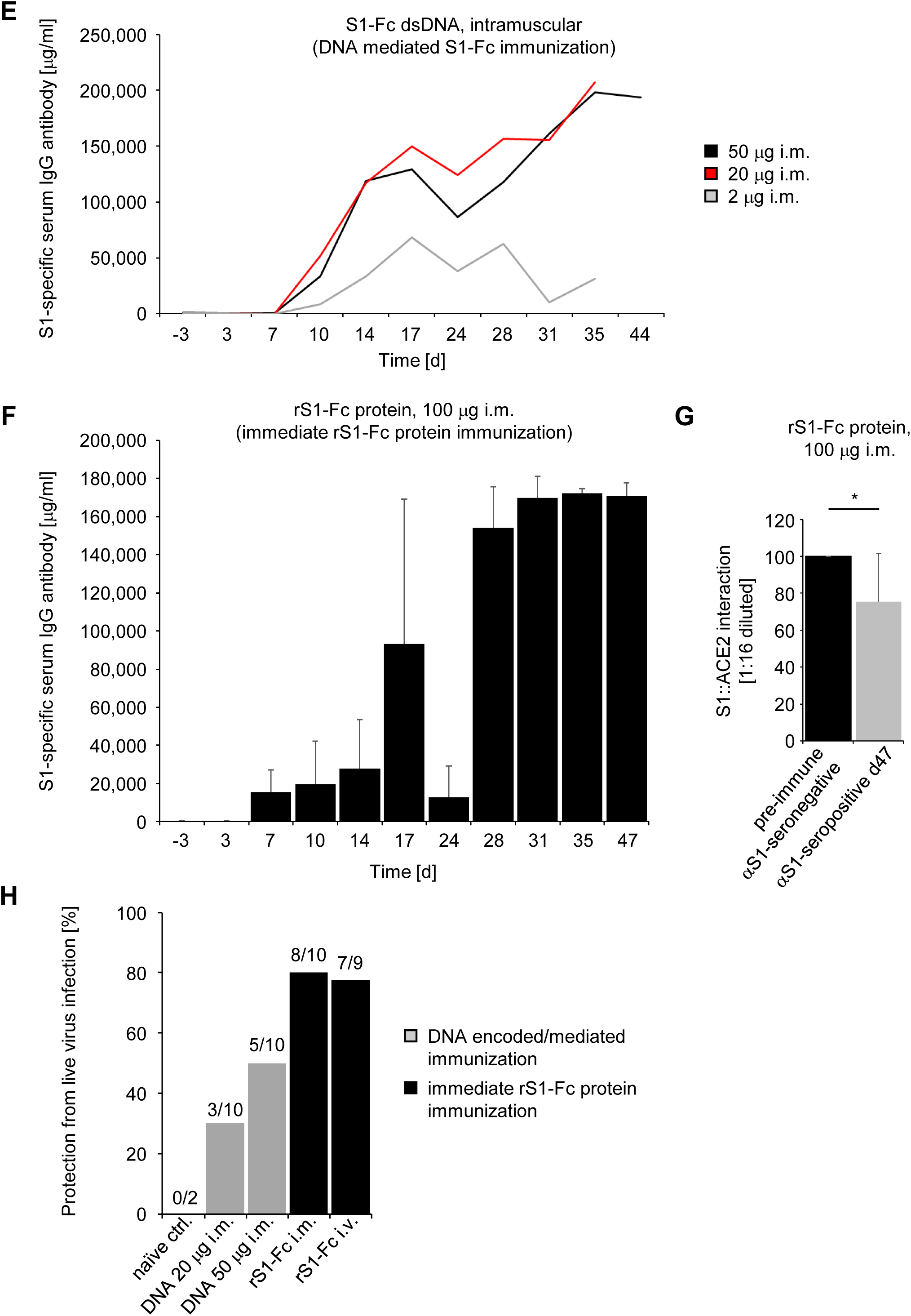
rS1-Fc immunization elicits early seroconversion, facilitating anti-S1-specific IgG production protecting against live SARS-CoV-2 challenge. (**A**) Schematic representation of mice immunization and blood serum collection schedule as indicated. (**B**) Dose-dependent and significant Th1 polarization upon high-dose non-expiring immunization with S1-Fc dsDNA as analyzed by fold-increased education of CD4^+^IFNγ^+^ versus CD4^+^IL-4^+^ T cells *in vivo* assessed by flow cytometry (right panel). Gating exemplary shown (left panels). (**C**) S1-antigen-specific CD8^+^ T cell accumulation assessed by MHC-tetramer/S1-peptide flowcytometric analysis of cognate TCR expressed by CD8^+^ T cells. (**D**) Immunization-dose-dependent maturation of effector CD8^+^IFNγ^+^ T cells was assessed by flow cytometry and quantified. (**E**) Rapid seroconversion and production of S1-specific IgG antibody upon immunization with S1-Fc dsDNA doses as indicated, or (**F**) rS1-Fc protein, assessed by ELISA. (**G**) Anti-S1 IgG seropositive blood serum significantly reduces viral S1:ACE2 host receptor interaction as shown by ELISA at high serum dilutions (1:16), and (**H**) elicits protective activity against live SARS-CoV-2 virus challenge, as assessed in a stringent experimental virus challenging assay employing VeroE6 cells. All *n*=10. SD shown; T-test: *) *P*<0.05.

## Discussion

Our results show that targeted immunization with S1-Fc immediately directed to the FcγR^+^ APC compartment induces a robust humoral immune response in preclinical testing eliciting protective activity against SARS-CoV-2 infection. S1-Fc immunization triggers rapid seroconversion, resulting in considerably elevated production of S1-specific antibodies *in vivo*. Furthermore, immunization with recombinant rS1-Fc protein improves the inhibitory activity of anti-S1 IgG antibodies, protecting from live SARS-Cov.2 infection *in vitro*. Interestingly, the education of an S1-antigen specific adaptive immune response is highly dose-sensitive. At high-dose immunization using non-expiring S1-Fc-encoding dsDNA, we demonstrate the desired Th1 polarization and effector CD8^+^IFNγ^+^ T cell maturation. Moreover, high-dose immunization with non-expiring S1-Fc-encoding dsDNA mounts S1-antigen specific CD8^+^ T cell education, which remained undetectable using low-dose immunization with rS1-Fc. Immunization dose-sensitivity shaping the adaptive immune response sheds light on potentially required elevated doses and/or an increased dose frequency for improved adaptive immune responses.

Impressive results were reported in a recent preclinical study, employing the full-length S glycoprotein of SARS-CoV-2 as immunogen, encoded and administered as plasmid DNA^26^, a conceptually very similar approach reported by Nabel and colleagues in 2004^11^. The bacterial plasmid DNA encoding full-length spike protein, regulated by a constitutively active cytomegalovirus (CMV) promoter and administered by electroporation, elicits an anti-SARS-CoV-2 but not anti-SARS or anti-MERS IFNγ T cell response, and the humoral immune response mounts serum IgG exerting neutralization activity upon a single administration of bacterial plasmid DNA encoding full-length spike protein^26^.

However, immunization introducing double-stranded, single-stranded DNA or RNA species including plasmids, inevitably raises the concern of an undesired Toll-like receptor (TLR) response, mediating uncontrolled cytokine secretion and potentially resulting in disease-promoting cytokine-release syndrome (CRS) and life-threatening inflammatory responses^27, 28^. TLRs are known to detect microbial infection extra-and intracellularly^29^, inducing the activation of an inflammatory response, including but not restricted to IL-6 production^30^. Moreover, TLR activation directly or indirectly compromises the functional priming of T cells by a blunted or indolent adaptive immune response, which *vice versa* insufficiently presents primary (MHC) and secondary (B7.1, B7.2) signals, resulting in propagation of anergic T cell populations^31^.

Moreover, although adenovirus-mediated introduction of immunogen, encoded by dsDNA inside the viral capsoid, aggressively achieves elevated antigen presentation over a considerably extended period of time, the systemic presence of adenovirus *per se* elicits unfavorable consequences^32, 33^. An undirected/anergic adaptive immune response, the education of inflammatory IL-10 producing CD4^+^ T regulatory cells reducing desired anti-viral CD8^+^ T cell efficacy, and raising undesired adverse reactions were reported^32, 33^, abandoning patient safety and is associated with unpredictable prognosis. Hence, the undirected and uncontrolled introduction of a transgene in humans for the purpose of immunization therapy must be considered scientifically premature, requires in depth clinical testing to determine safety exit strategies, and is not fit for delivery to the public at large scales, which are required to control the COVID-19 pandemic.

Our ongoing comprehensive studies as well as others^26, 32, 33^ indicate, that considerable amounts of immunogen are required to achieve education of an IFN^+^ adaptive immune response. However, low-dose immunization with rS1-Fc targeting the FcγR^+^ APC compartment represents an operative immunization strategy achieving rapid seroconversion and mounting considerably high levels of anti-S1 specific antibodies eliciting protective activity. Further preclinical studies tailored to improve the antigen-specific adaptive immune response in the circuitry of immunization by targeted rS1-Fc protein are required, guiding us to a mild, safe and protecting targeted rS1-Fc vaccination against COVID-19.

## Material and methods

### Mice and cell culture

For intramuscular antigen challenge, C57BL/6 female mice 5-6 weeks old were injected with either 100 μg rS1-Fc, or electroporation was applied delivering 2 μg, 20 μg, or 50 μg linear dsDNA encoding S1-Fc. For intramuscular antigen challenge, C57BL/6 female mice were injected with 100 μg rS1-Fc intravenously. Antigen challenge at day 1 and day 22 was administered into the *biceps femoris* muscles of mice. Peripheral blood was collected from anaesthetized mice once/week via retro-orbital route. Murine NIH/3T3 fibroblasts, C2C12 myoblasts and 264.7 RAW macrophages were obtained from ATCC and subcultured according to the vendor’s instructions. Single-cell suspensions from spleens were freshly isolated immediately after dissection. Briefly, spleens were disrupted using cell strainer, followed by red cell depletion.

### Production of linearized S1-Fc-encoding S1-Fc dsDNA fragment

The plasmid pAAV S1-Fc was digested with New England Biolabs (NEB) Restriction Enzymes Pvu I-HF (Cat. No. R3150), Sph I-HF (Cat. No. R3182) and Mlu I-HF (Cat. No. R3198) at 37°C for 2.5 hours. The resulted fragments were separated by 1% agarose DNA gel electrophoresis. The desired S1-Fc fragment was cut from the gel and extracted to recover S1-Fc DNA with NucleoSpin Gel and PCR Clean-up kit from Macherey Nagel (Cat. No. 740609). The S1-Fc dsDNA was amplified by 2-step nested PCR. The gel recovered S1-Fc DNA was used as the template for the 1st PCR of the nested PCR. Both 1st PCR and 2nd PCR were performed by using PrimeSTAR HS DNA polymerase from Takara Bio. (Cat.No. R010A). The large scale of PCR was performed in multiple 96-well PCR plates. In each well, 50 μl PCR reaction solution contains 1X PCR reaction buffer, 0.2 mM dNTP, 20 ng template, 0.4 μM of each primer, and 1.25 units of polymerase. The PCR product was purified with anion exchange column AX2000 (Macherey-Nagel, Cat. No. 740525) and precipitated by IPA (IBI Scientific, Cat.No. IB15730). The DNA pellet was washed with 70% ethanol (VWR, Cat. No. 71006-012) twice and air dried. The DNA pellet of dsDNA S1-Fc was dissolved in PBS buffer (Corning Cat.No.21-040-CM) at 2-2.5 μg/μl. Qubit dsDNA Broad Range Assay Kit (Thermo Fisher Scientific, Cat. No. Q32850) was used for dsDNA S1-Fc concentration determination. The identity and purity of dsDNA S1-Fc was verified by DNA gel electrophoresis imaging.

### Expression of recombinant rS1-Fc protein

rS1-Fc was expressed in Chinese hamster ovary cells (Free Style CHO-S Cells by Thermo Fisher Scientific, # R80007) in CHO-S-SFM media with hypoxanthine and thymidine (Life Technologies, # 12052098) using polyethylenimine (PEI) for DNA delivery at a ratio of 4:1. Transfection was performed at a cell density of 2 × 10^6^ cells/ml at 37°C for 24 hours. The addition of CD FortiCHO Medium (Life Technologies, # A1148301) with 20% Penicillin Streptomycin: Amphotericin B solution Cell Culture Reagent 100X (# 091674049) and 100X GlutaMax supplement by Gibco (# 35050061) was added the following day and transferred to 28°C. Viability was monitored by Contessa II (Invitrogen) and antibody concentrations were monitored via Octet. The rS1-Fc was protein A purified using MabSelect SuReLX resin (# 1754702) obtained from GE Healthcare.

### Fluorescent labeling of rS1-Fc

rS1-Fc Antigen (STI) was conjugated to IRDye® 800RS NHS Ester (Li-cor) using amine-reactive crosslinker chemistry. Briefly, S1-Fc antigen (2.5-3.0 mg/ml) was reacted with 5 equivalents of IRDye® 800RS NHS Ester in 1x DPBS pH 7.4 containing 5% anhydrous DMSO for 3 hours with gentle rotation at room temperature. Subsequently, the reaction mixture was subjected to PD-Minitrap G-25 column (GE Healthcare) to remove unreacted dyes according to the manufacturer’s instructions. Upon purification, the conjugate underwent buffer exchange three times into 1x DPBS pH 7.4 using a 4-ml Amicon Ultra centrifugal filter (30 kDa MWCO, Millipore). The conjugate was characterized using SDS-PAGE, SEC HPLC, and BCA Assay.

### Longitudinal near infrared in vivo imaging of lymphatics

IRDye800-rS1-Fc was injected at 10 μl (1 mg/ml) intramuscularly into the *biceps femoris* of C57BL/6 mice. Near-infrared fluorescence imaging kinetics at 100 ms exposure time was performed upon administration. The injection site was concealed to prevent photonic overexposure. Fluorescent rS1-Fc homing to/clearing from the inguinal lymph node was quantified by applying a region of interest (ROI) to determine fluorescent intensity kinetics.

### Detection of anti-SARS-CoV-2 Spike S1-domain specific serum antibody

A direct binding ELISA format was used to detect the anti-SARS-CoV-2 Spike S1 Subunit antibody in mouse serum samples. Plate was coated with SARS-CoV-2 (2019-nCoV) Spike Protein (S1 Subunit, His Tag; Sino Biological, Cat# 40591-V08H) at 5 μg/ml at 4°C overnight. Next day the plate was washed 3 times with 1x KPL buffer (Sera Care, Cat# 5150-0008) and blocked in Casein Block Buffer (Thermo Scientific, Cat# 37528) at room temperature (RT) for 1 hour. A mouse monoclonal antibody (SARS-CoV Spike S1 Subunit Antibody) from Sino Biological (Cat#40150-MM02) was used to determine the standard curve between 100 – 0.781 μg/ml. The standard was prepared using Casein Block Buffer at a 2-fold serial dilution. Mouse serum samples were diluted with Casein Block Buffer at the designated dilution factor during plate blocking period. The blocked plate was washed once and incubated with the standard or test samples at room temperature for 1.5 hours with shaking at 300-400 rpm. The plate was washed 3 times prior to adding 50 μl of 1:1000 diluted Goat anti-Mouse IgG H+L-HRP antibody (Bio-Rad, Cat# 172-1011) to the plate and incubated for 1 h at RT, 300 rpm. The plate was washed for 3 times prior to adding 50 μl of TMB substrate (Thermo Scientific, Cat# 34021) to each well. The plate was incubated at RT with shaking for 10 minutes. At end of incubation, 50 μl of 2 M Sulfuric Acid was added to each well to stop TMB development and optical density OD was assessed at λ=450 nm immediately. Sample concentration (μg/ml) was determined by fitting the tested sample data, *i*.*e*. Y= A450 Absorbance with blank subtracted, to a 4-parameter logistic curve generated by the standard serials using non-linear regression in SoftMax Pro GxP.

### Tissue Section Indirect Immunofluorescent Staining and Confocal Microscopy

Collected tissue frozen in O.C.T. media (Fisher Scientific) was dehydrated and sections underwent immunostaining procedure as previously described^34^. Briefly, tissues were fixed with 2% paraformaldehyde and permeabilized with icecold methanol for 30 min gently shaking. Wax circles were applied and Image IT signal enhancer (Invitrogen) was used for signal quenching. Upon blocking tissue with PBS containing 10 % mouse serum and 2.5% goat serum for 1 h at RT, primary antibodies against S1 or dendritic cell marker (clone 33D1) were incubated diluted in blocking buffer in a wet chamber over night at 4°C. Upon rinsing, fluorescent secondary antibodies diluted in blocking buffer were incubated together with DNA staining DRAQ7 (abcam) for 4 hours protected from light and sections were mounted in Mowiol (Sigma-Aldrich). Stained sections were analyzed by confocal laser-scanning microscopy using a cLSM510Meta Microscope (Zeiss).

### Intracellular Staining and Flow Cytometry

To prepare single-cell suspensions for flow cytometry, spleens from naïve or vaccinated mice were collected and homogenized in PBS^35^. Digests were filtered through 70 μm cell strainers, centrifuged at 1,500 rpm for 5 min. After red blood cell lysis (Sigma-Aldrich), single-cell suspensions were filtered, washed, and resuspended in FACS Wash Buffer (2% FBS in Hank’s balanced salt solution without Ca, Mg, and phenol red). Single-cell suspensions were stimulated for 3 hours with PMA (5 ng/ml, Sigma) and inomycin (500 ng/ml, Sigma) in the presence of protein transport inhibitor (monensin 1000x, Biolegend). For antigen-specific study, single-cell suspensions were stained with MHC-tetramer mounted with spike protein peptides (MBL international). Cells were blocked with CD16/CD32 and incubated for 30 min on ice with FITC-, PE-, APC-, PECy7-, PerCP-Cy5.5, BV421, BV605, BV 650, and BV785-conjugated surface-staining antibodies (CD3, CD4, CD8, CD69, CD19, CD11c, CD80, CD86, and F4/80), followed by intracellular staining antibodies (IL-4, IFNγ, and S1) 30 min on ice, purchased from Biolegend, or in-house. Aqua LIVE/DEAD used for cell viability was purchased from Invitrogen. Cells were washed twice before analysis on the BD FACSCelesta flow cytometer (Becton Dickinson).

### Meso Scale Discovery MSD ELISA

Wells of an MSD plate were coated with hACE2 protein (2 μg/ml in PBS) over night at 4°C. After coating, the hACE2 solution was removed followed by blocking with Superblock (Scy Tek Labs Cat#AAA500, Lot# 55263) for 1 h on a horizontal shaker (100 rpm). Subsequently, the wells were washed with KPL buffer (Sera Care Cat#5150-0009, Lot#10408687) and cell culture media supernatant of electroporated C2C12 cells was added in triplets with a dilution of 1:5 in Superblock, starting with the undiluted supernatant. S1 protein was used as a standard (starting concentration 40 μg/ml, 1:5 dilution). The plate was incubated at RT for 2 h on a horizontal shaker (100 rpm). After washing with KPL buffer, the wells were incubated with αS1 antibody at a concentration of 1 μg/ml in Superblock. Upon washing with KPL buffer, the wells were incubated for 2 h with the sulfo-tagged antihuman IgG (MSD Cat# R32AJ-1, Lot# W0015871S) in a dilution of 1:1000 in Diluent 100 (MSD Cat#R50AA-2, lot# 0270266). After the incubation, the wells were washed and 1x Read buffer (MSD Cat# R92TC-2, Lot#Y0140370) was added. The OD assessment was carried out by the MSD QuickPlex SQ 120. The data was analyzed via the included software.

### Virus neutralization assay

Serum samples from vaccinated mice were heat-inactivated with 56°C for 30 minutes. Equal amounts of 2-fold diluted serum samples and diluted SARS-CoV-2 (2019-nCoV/USA-WA1-A12/2020) yielded to 100 TCID50/50 µl were mixed and incubated at 37°C for 30 minutes. Confluent Vero E6 cells in 96-well plates were inoculated with 100 µl of the virus-serum mixture. After 1 hour absorption, the cells were washed and culture media was replaced. The cells were fixed with 10% formalin and stained with crystal violet at 72 hours post-inoculation to check the cytopathic effect. The serum samples which had neutralizing activity at 1:2 dilution were evaluated as neutralization positive.

### Study approval

Mouse care and experimental procedures with mice were performed under pathogen-free conditions in accordance with established institutional guidance and approved Animal Care and Use Protocols (ACUP) from the Research Animal Care Committee at Sorrento Therapeutics Inc.

## Acknowledgments and contributions

We thank our staff at Sorrento Therapeutics Inc. to tirelessly support our efforts.

HJ drafted the study design, orchestrated the study, and edited the manuscript, AH aligned required experimental procedures, performed and analyzed experiments, and wrote the manuscript. CY, TA and CL performed and analyzed experiments and contributed to manuscript writing. HZ, WG, LK and YZ designed and produced the immunogen and contributed to manuscript writing. WS, SY and SK performed ELISA assays and nIR imaging and contributed to manuscript writing. YF produced fluorescently labeled immunogen. RA and MB contributed to manuscript editing and scientific discussion. SP and JM performed and analyzed experiments using live SARS-CoV-2 virus and contributed to manuscript edition.

## Additional Information

Correspondence and other requests should be addressed to Andreas Herrmann, Slobodan Paessler, and Henry Ji.

