## Supplementary figures for "A Targeted Vaccine against COVID-19: S1-Fc Vaccine Targeting the Antigen-Presenting Cell Compartment Elicits Protection against SARS-CoV-2 Infection"

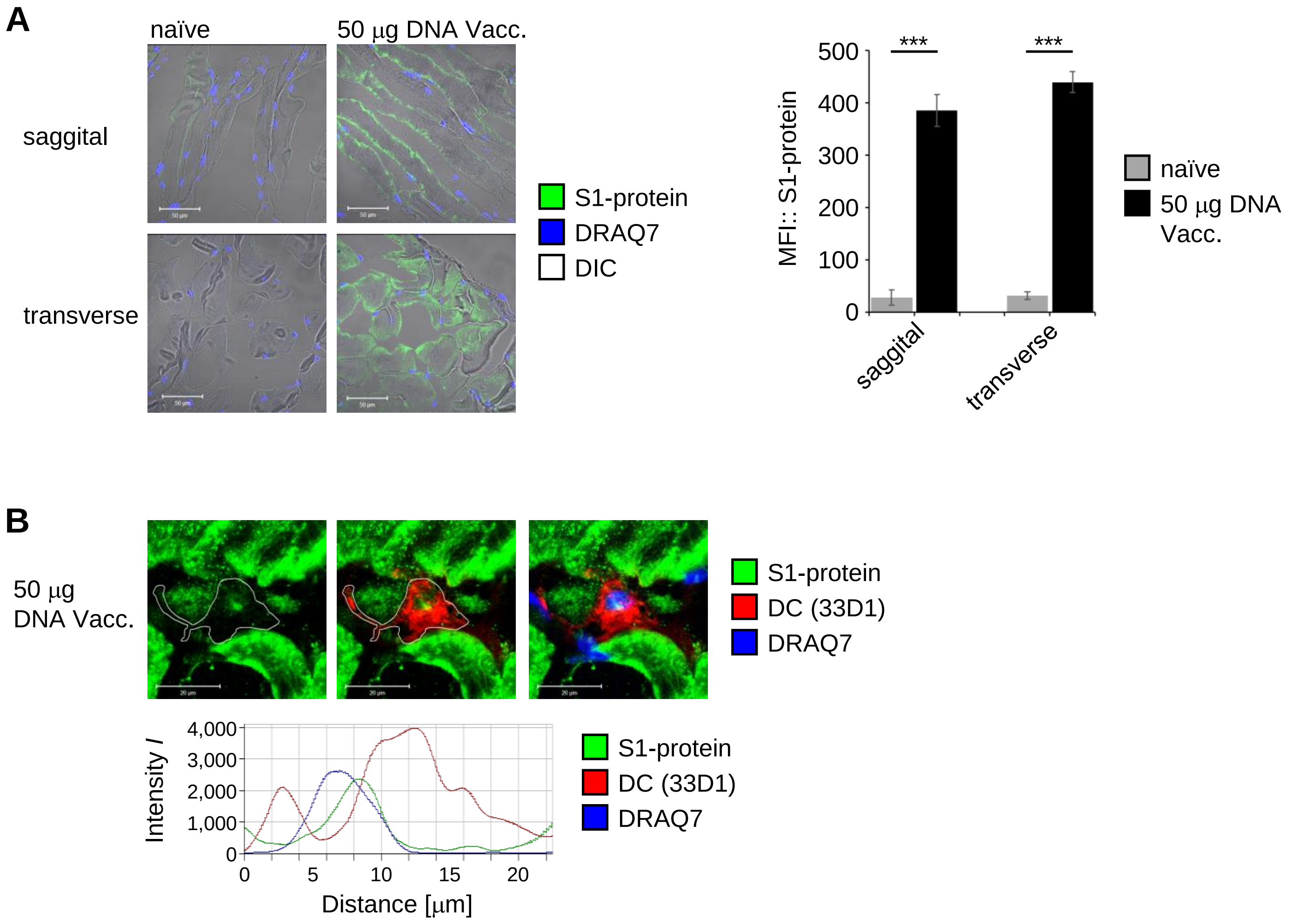


**Supplemental Fig. 1: S1-Fc protein expression in murine *biceps femoris* and uptake by dendritic cells *in vivo*.** Upon immunization with non-expiring S1-Fc encoding dsDNA, (**A**) S1-Fc (green) expression in murine muscle *biceps femoris* was assessed on day 47 by confocal microscopy and quantified. SD shown, T-test: ***) P<0.001. Scale, 50 μm. (**B**) S1-protein (green) produced *in vivo* by immunization with non-expiring S1-Fc encoding dsDNA, was internalized by dendritic cell (red). A fluorescence profile illustrates S1 cellular uptake (green within red cell borders, blue nucleus). Scale, 20 μm.

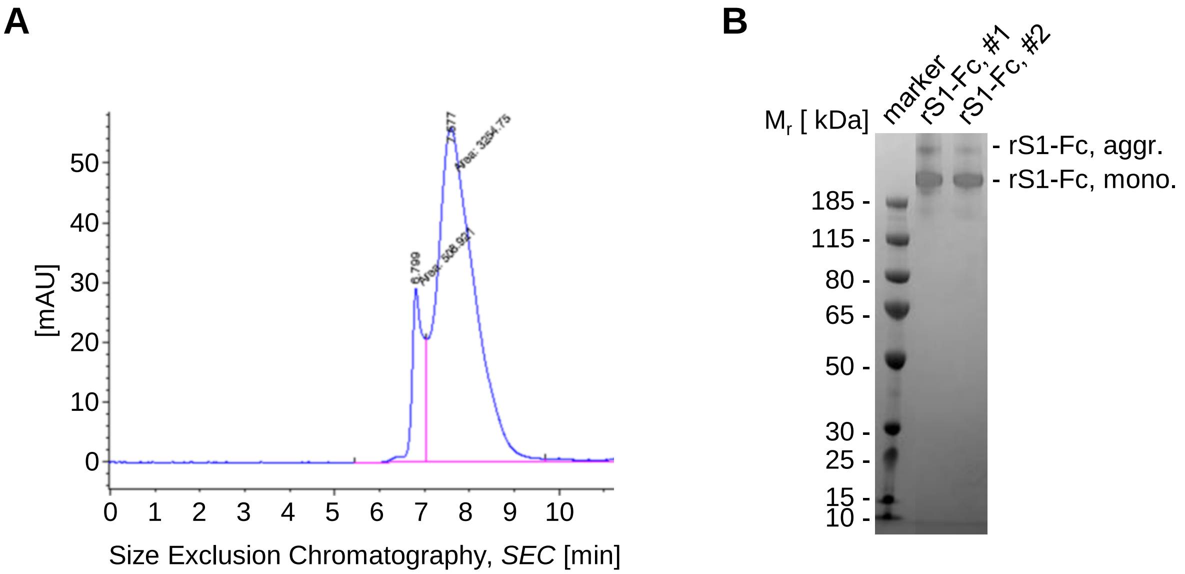


**Supplemental Fig. 2: rS1-Fc protein production at Sorrento Therapeutics.** (**A**) Recombinant rS1-Fc was expressed in CHO cells and produced rS1-Fc was analyzed by SEC HPLC to assess monomer purity and intrinsic oligomerization aggregation. (B) Two independent batches of produced rS1-Fc were subjected to electrophoretic protein separation and visualized by *Coomassie* protein staining.
